# Rapid and robust antibody Fab fragment crystallization utilizing edge-to-edge beta-sheet packing

**DOI:** 10.1101/2020.04.14.040949

**Authors:** Ricky Lieu, Stephen Antonysamy, Zhanna Druzina, Carolyn Ho, Rebecca Kang, Anna Pustilnik, Jing Wang, Shane Atwell

**Affiliations:** Biotechnology Discovery Research, Applied Molecular Evolution, Eli Lilly and Company; San Diego, CA, USA; Discovery Chemistry Research and Technologies, Eli Lilly and Company Corporate Center, Indianapolis, IN, USA

**Author notes:** Corresponding author (SA). Department of Biomedical Engineering, Boston University, Boston, MA, USA.

## Abstract

Antibody therapeutics are one of the most important classes of drugs. Antibody structures have become an integral part of predicting the behavior of potential therapeutics, either directly or as the basis of modeling. Structures of Fab:antigen complexes have even greater value. While the crystallization and structure determination of Fabs is easy relative to many other protein classes, especially membrane proteins, broad screening and optimization of crystalline hits is still necessary. Through a comprehensive review of rabbit Fab crystal contacts and their incompatibility with human Fab structures, we identified a small secondary structural element from the rabbit light chain constant domain potentially responsible for hindering the crystallization of human Fabs. Upon replacing the human kappa constant domain FG loop (HQGLSSP) with the two residue shorter rabbit loop (QGTTS), we dramatically improved the crystallization of human Fabs and Fab:antigen complexes. Our design, which we call “Crystal Kappa”, enables rapid crystallization of human fabs and fab complexes in a broad range of conditions, with less material in smaller screens or from dilute solutions.

## Introduction

Antibody therapeutics are one of the most important classes of drugs. By the end of 2019, 90 monoclonal antibody drugs covering immune disease, infection disease, cardiovascular disease, cancer and others had been approved in the U.S. and Europe [1], accounting for a projected $150B [2]. While at one time rodent antibodies were developed for human use, this was followed by a long period of humanized antibodies, which over the last two decades has shifted to entirely human discovery platforms like phage [3] and yeast display [4] or by immunization of rodents with human germline repertoires [5]. In these platforms engineering is not necessary for humanization but continues to be used to address other issues: affinity, cross-reactivity, post translational modifications, hydrophobicity, electrostatics, viscosity, and immunogenicity. Furthermore, characterization of antibodies continues to become more sophisticated, especially as new antibody derived formats are developed like antibody drug conjugates and bispecific antibodies [6].

Modeling of antibody structures has become an integral part of predicting the behavior of potential therapeutics, especially for properties such as hydrophobicity, stability, charge/dipole moments and deamidation propensity [7]. This modeling is typically based on the publicly available crystal structures with the most similar CDR sequences. Due to the difficulty of modeling CDRs, especially heavy chain CDR3, calculations based on the crystal structures of the exact (or highly similar) Fab crystal structures should improve the accuracy of antibody property predictions.

Structures of Fab:antigen complexes have even greater value. They can supply the crystal structure of the Fab for the above and also epitope:paratope information. Obtaining the Fab:antigen complex structures is the only way to directly determine the relative 3-dimensional positions of the antigen and Fab to the precision of individual atoms. Epitopes can be seen, rather than inferred. Amino acid side chains can be examined and hypotheses formed regarding their roles in affinity and cross-reactivity and calculations conducted to predict and engineer affinity (up or down) and cross-reactivity. In addition, the structure of a complex can be referenced when considering other mutations and can be an essential cross-check in determining the validity of different assay formats yielding confusing or contradictory results.

However, crystal structure determination is challenging and costly [8]. Estimates from a decade ago put the all-in cost around $50,000 [9]. The most difficult step tends to be the production of well-ordered crystals from purified protein. There are many factors that may hinder the crystallizability of a protein: purity, stability, disorder (inter domain, loop or termini), surface charge and hydrophobicity, etc. [10]. Obtaining well diffracting crystals can take a few days or a few years or might simply be abandoned after significant effort.

While the crystallization and structure determination of Fabs is easy relative to many other protein classes, especially membrane proteins, broad screening and optimization of crystalline hits is still necessary. Like other proteins some Fabs require significant optimization and examples of entirely recalcitrant Fabs exist. Fab:antigen complexes are often easier to crystallize than the antigen alone (hence the use of Fabs as “crystallization chaperones” [11]) but can still be difficult and require extensive screening and optimization. The individual attention across days to months required in crystallization and structure refinement make these two steps the most expensive in the process from sequence to final structure. Difficult cases are especially and negatively impactful to overall averages. The effort required for Fab crystallization perhaps explains why so few structures have been used for engineering or calculations.

By engineering a small secondary structural element from the rabbit light chain constant domain into the human kappa light chain, we dramatically improved the crystallization of human Fabs and Fab:antigen complexes.

## Results

### Analysis of rabbit Fab crystal packing and design of human crystallizable kappa

Visual inspection of the crystal packing interactions in 36 published and Lilly proprietary rabbit Fab crystal structures showed that constant domain beta-strand to beta-strand crystal packing was common (Table 1). In 68% of the 19 deposited rabbit Fab structures (including Fab complexes) a LC to LC beta interaction occurs, forming a continuous beta sheet across two Fab molecules (Fig 1A). Over a third of those structures also have a HC:HC beta packing interaction. Overall, 84% of the structures form some kind of beta sheet packing interaction in the G strand of the constant domains.

**Table 1.** G-strand beta sheet crystal packing of rabbit Fab and Fab complex structures. Percentage of deposited PDB structures with either HC:HC, LC:LC or HC:LC G-strand beta sheet formation. Also indicated are percentage with both HC:HC and LC:LC packing as well as percentage of structures with any G-strand beta packing. Lilly internal statistics are listed below PDB statistics.

| **PDB** | **Res** | **HC:HC** | **LC:LC** | **HC:LC** |
| --- | --- | --- | --- | --- |
| **4HBC** | 1.54 |  | x |  |
| **4JO1** | 2.03 |  |  | x |
| **4JO2** | 2.50 |  |  | x |
| **4JO3** | 2.60 | x | x |  |
| **4JO4** | 2.27 | x | x |  |
| **4ZTO** | 2.30 | x | x |  |
| **4ZTP** | 1.63 |  | x |  |
| **5DRN** | 1.99 |  | x |  |
| **5DS8** | 1.95 |  | x |  |
| **5DSC** | 2.40 |  | x |  |
| **5DTF** | 1.90 | x |  |  |
| **5DUB** | 2.00 |  |  |  |
| **5M63** | 2.74 | x | x |  |
| **5V6L** | 2.55 | x | x |  |
| **6CEZ** | 2.40 |  |  |  |
| **6CJK** | 1.80 |  | x |  |
| **6I9I** | 1.98 |  |  | x |
| **6PEH** | 2.30 | x | x |  |
| **6T3F** | 3.20 |  | x |  |
| **PDB Total** | **19** | **37%** | **68%** | **16%** |
| **HC:HC & LC:LC** |  | **32%** |  |  |
| **Any** |  | **84%** |  |  |
| **Lilly Internal** | **17** | **65%** | **71%** | **6%** |
| **HC:HC & LC:LC** |  | **59%** |  |  |
| **Any** |  | **82%** |  |  |

**Fig 1.**
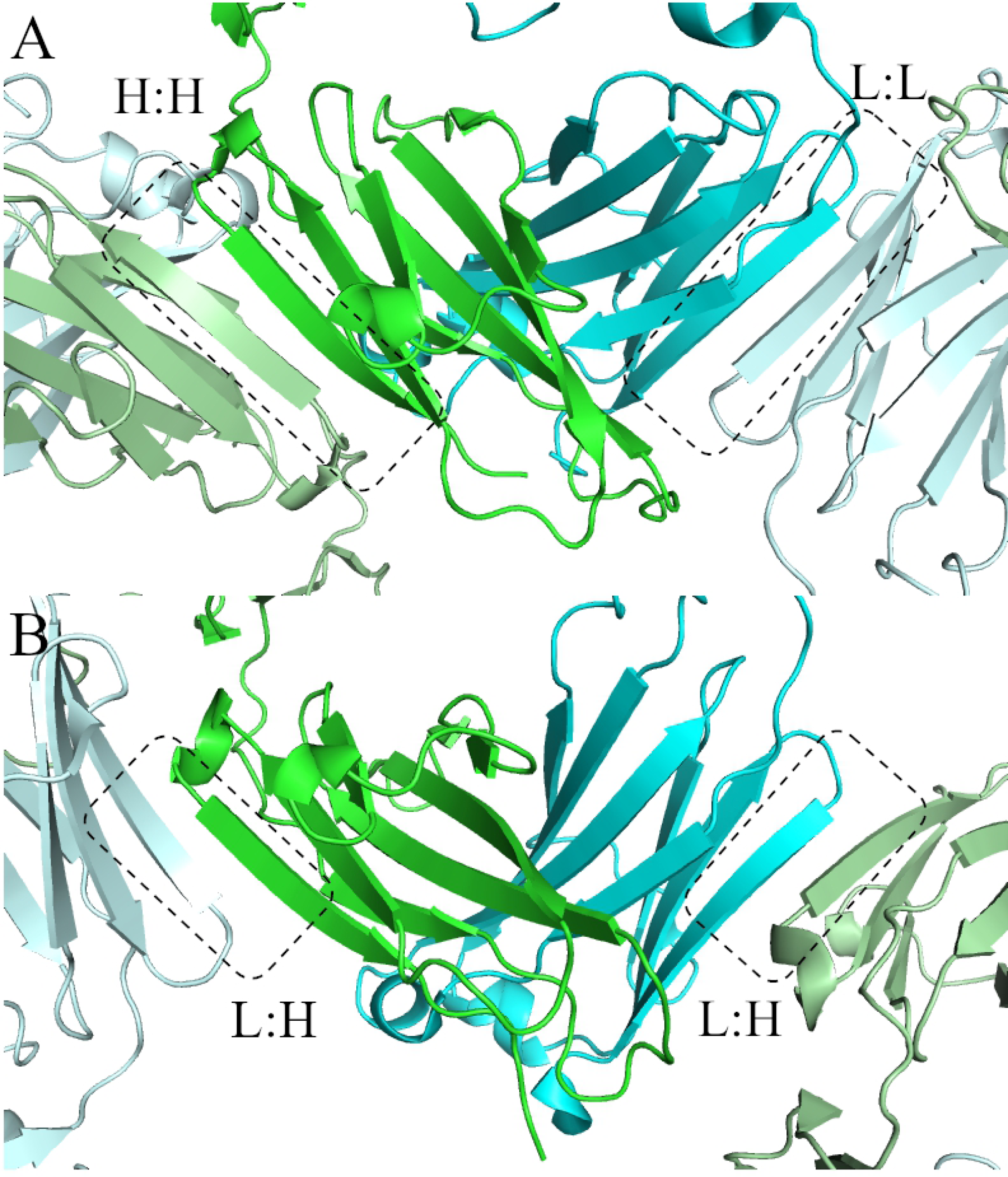
Examples of G-strand beta sheet packing in rabbit Fab crystal structures. (A) HC:HC and LC:LC beta packing in 4ZTO.pdb. Heavy chain in green and pale green. Light chain in Cyan and pale cyan. (B) HC:LC beta packing in 4JO1.pdb. Figures prepared in Pymol.

Our in-house experience is similar but with more examples of Fabs that have both HC:HC and LC:LC crystal packing interactions and fewer examples that have HC:LC packing (Table 1 & Fig 1B). Crystals of Fabs that form both the HC:HC and LC:LC or that form HC:LC packing interactions have a continuous column of constant domains, each domain forming a typical Fab HC:LC interaction as well as a beta sheet with another domain.

A survey of dozens of human Fab structures showed no such interactions. Alignment of a human Fab structure onto a rabbit Fab showed that the longer FG loop present in human Kappa constant domains forms a bent and bulging conformation that would interfere with a beta sheet packing interaction (Fig 2A). This longer FG loop is shared by mouse Kappa constants but not by rabbit Kappa constants (nor by lambda constant domains) (Fig 2B). Visual inspection of the human Fab aligned on the rabbit packing interaction also suggested that the human kappa constant Lysine 126 (Kabat numbering) might be an impediment to beta sheet packing on the opposite side of the domain (Fig 2C).

**Fig 2.**
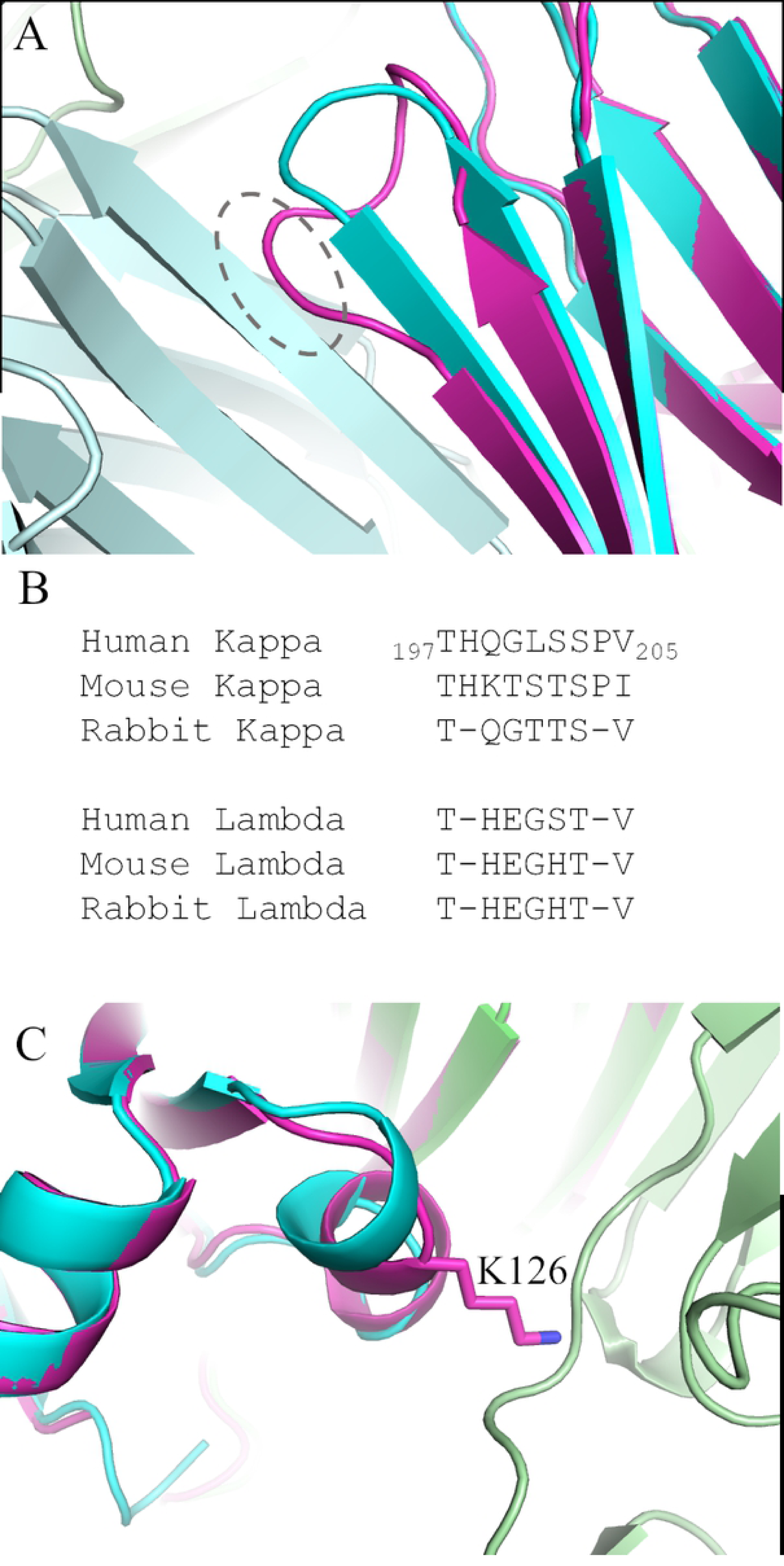
Incompatibility of human kappa constant FG loop with rabbit like LC:LC packing. (A) Structural alignment of the human fab from 4NZU.pdb (magenta) on the rabbit Fab crystal packing from 4ZTO (cyan and pale cyan) shows that the rabbit Constant Kappa FG loop (cyan) is much more compact and compatible with the beta sheet packing and that the longer and bulging human loop (magenta in dashed grey circle) would not be. (B) Sequence alignment of FG loops of human, mouse and rabbit kappa constant domains and lambda constant domains. Rabbit loops are two residues shorter and lack a proline. They resemble the FG loop from the lambda constant domains. (C) Second potential site of interference is the human K126 sidechain. Alignment shows that it could interfere with HC:HC packing.

We designed several variants of a hexahistidine tagged Fab fragment to test the impact of replacing the human sequences (either the FG loop or K126) with their rabbit equivalents. In the case of the FG loop, this consisted in replacing the septamer human sequence HQGLSSP (positions 198 to 204 according to Kabat numbering) between the structurally homologous T and V (position 197 and 205, respectively, according to Kabat numbering), with the pentamer rabbit sequence QGTTS (positions 199 to 203 according to Kabat numbering). The resulting mutant is shorter by two residues, with deletions of histidine at position 198 and proline at position 205, and is referred to here as ΔQGTTSΔ (positions 198 to 204 according to Kabat numbering). In several variants, Lysine 126 was mutated to Alanine (K126A, Kabat numbering).

Separately to the crystal packing analysis we have observed that the C-terminal interchain disulfide is rarely ordered in Fab structures. This could be due to conformational heterogeneity or to heterogeneous oxidation. To address the later we designed two variants: one (“GEP*”) with the disulfide simply removed by mutating the C-terminal kappa chain cysteine to proline (position 214 according to Kabat numbering) and the (IgG4) Cys127 to alanine (C127A, Kabat numbering); the second design (“ESKCGGH6”) mutated the kappa cysteine to proline and created a new disulfide partner at the C-terminus of the heavy chain by mutating Tyr 229 to Cysteine.

### Crystallization of Fabs

The results of incorporating these mutations singly and in combination into two Fabs and attempting crystallization is shown in Table 2. We incorporated the described mutations into two Fabs. One is derived from the publicly available dupilumab (Dupixent ™) sequence. The second was the part of an internal discovery effort against a cell surface receptor. Two 96-well crystallization screens were used with Fabs purified identically, set up at approximately 10 mg/ml, streak seeded with unrelated Fab crystals (in order to eliminate stochastic differences due to nucleation) and analyzed at the same time point (9 days). Both parental Fabs produced crystalline hits in a few conditions and the dupilumab Fab crystals even yielded a 2.0Å dataset and structure. Neither the K126A mutation (Table 2) nor any of the disulfide variants (not shown) produced significantly more conditions with crystals. The most dramatic difference was upon incorporating the ΔQGTTSΔ FG loop. The Fabs with this loop yielded crystals in approximately 100 conditions out of 192 for the G6 Fab and 125 out of 192 for the dupilumab Fab. Crystals harvested directly out of these screens (i.e. not optimized in subsequent screens) produced high resolution datasets (Table 2). The best dupilumab Fab diffracted to 1.4Å from the ΔQGTTSΔ alone and the best G6 Fab to 1.15Å from a combination of the FG loop mutation with K126A and the intrachain disulfide.

**Table 2.**
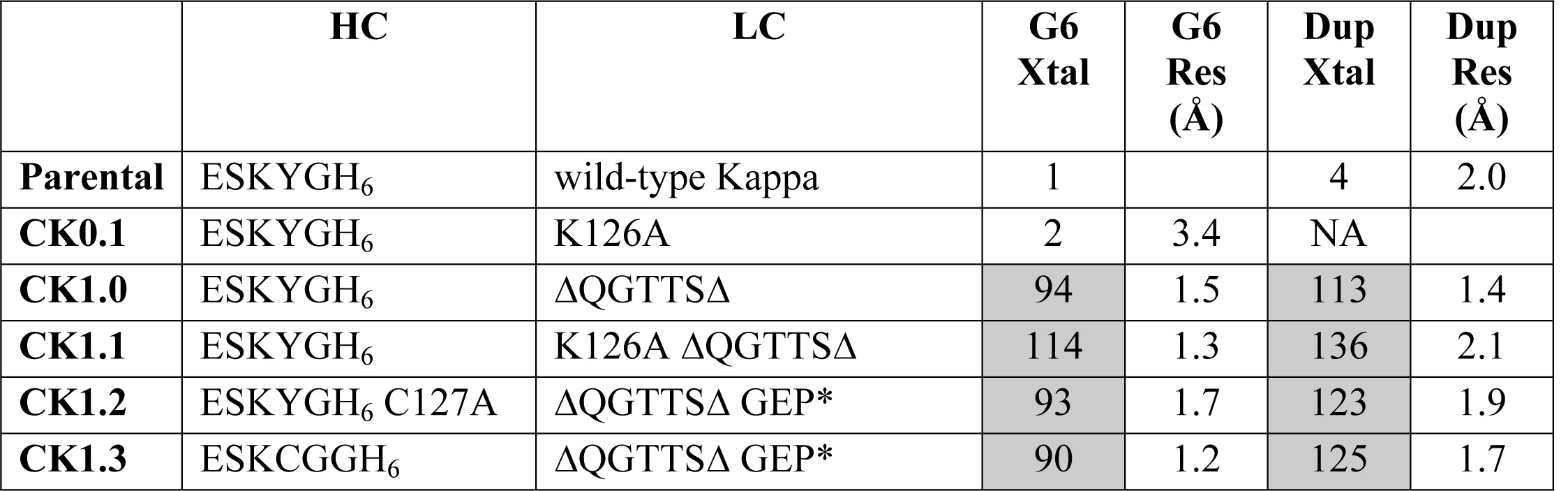
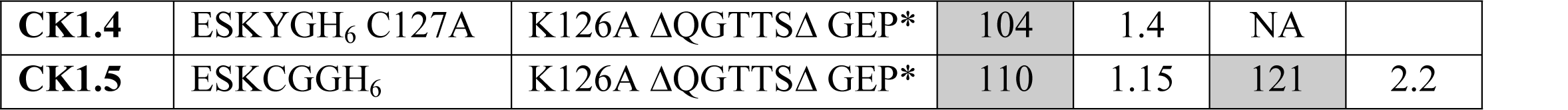
Crystallization of two fabs and their variants. The parental light chain was the wild-type kappa. The parental heavy chain terminated at the sequence ESKYG and included six histidines for purification purposes. The ‘Xtal’ column indicates how many conditions from two 96-well screens produced any kind of crystal by approximately day 9. Several crystals were sent from each construct with harvestable crystals and the best resolution dataset is indicated.

### Crystal structures of engineered Fabs

Fifty-nine datasets were collected for the G6 variants, encompassing 11 crystal forms. Structures were solved for 7 and refined for 5 crystal forms: P2_1_2_1_2_1_ with a 43×75×165 Å cell (5 refined structures), P4_3_2_1_2 77×77×330 (3), P2_1_2_1_2_1_ 66×74×91 (1), P1 53×65×67 85×71×84 (1), and C2 206×103×70 β=92.7° (1). Forty-four datasets were collected for the dupilumab Fab variants, encompassing 11 crystal forms. Structures were solved for 4 and refined for 3: P2_1_ 53×66×135 β=91.6° (1), P2_1_2_1_2_1_ 59×73×105 (1), P4_3_2_1_2 74×74×185 (1).

Structures of the Fabs variants without the ΔQGTTSΔ, including the parental, did not pack with extended beta sheet interactions. The parental dupilumab Fab for example packs with various types of interactions, but none that form a continuous beta-sheet (Fig 3A). All structures derived from Fabs with the ΔQGTTSΔ FG loop on the other hand packed forming a beta sheet between the G strand of the kappa constant domain and the G strand of the constant heavy 1 (CH1) domain (Figs 3B and C). This interaction is similar but not identical to that was seen in rabbit Fab crystal packing. It involves the same strands as seen in Fig 1B, but is more extensive, involving 7 residues on both sides like the H:H or L:L interactions seen in Fig 1A. The pseudo-symmetric center of this beta-sheet is between kappa V208 (position 205 according to Kabat numbering) and CH1 V219 (Kabat numbering).

**Fig 3.**
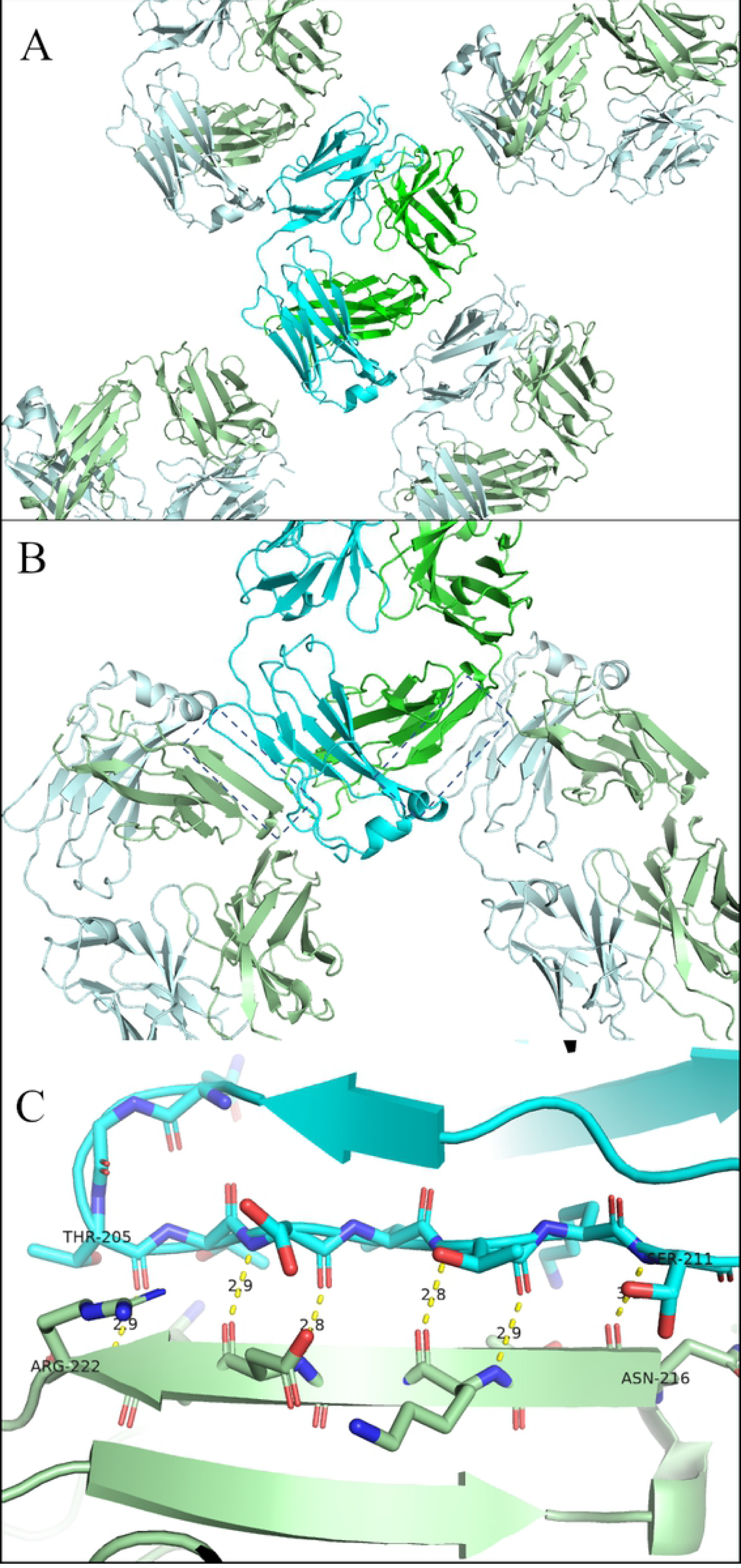
Crystal packing in CK constructs. (A) Crystal packing in un-engineered dupilumab parental Fab. No G strand beta packing is present. (B) One plane of crystal packing in crystal kappa version. Each kappa constant (blue and light blue) forms a beta sheet with a nearby constant heavy 1 domain (green and pale green). (C) beta sheet formed between the G strand of the constant kappa domain (cyan) and the G strand of the constant heavy 1 domain (pale green). Sheet extends from kappa T205 to S211 and heavy N216 to R222 which is pseudo-symmetric and centered between the kappa V208 and CH1 V219.

The K126A mutation does not appear to impact crystal packing or diffraction quality. The highest resolution structure for the G6 Fab did incorporate this mutation, but the impact of the mutation was not systematic. The highest resolution structure for the dupilumab Fab for example incorporated only the FG loop mutation ΔQGTTSΔ. Nor did the disulfide removal (C127A + GEP*) or intrachain disulfide (ESKC + GEP*) appear to impact crystal packing or diffraction. Structures were obtained with the intrachain disulfide ordered. The temperature factors in this region were higher than average as is seen in other structures with the interchain disulfide ordered.

### Crystallization of Fab:antigen complexes

We next applied the crystallization mutations to Fab:antigen complexes. In the case of G6, we utilized the CK1.5 variant of the Fab since it had diffracted the best as a Fab alone. For dupilumab Fab and the other 4 complexes, we utilized the simpler CK1.0 (i.e. only ΔQGTTSΔ). All CH1s were IgG4 and had the same C-terminal hexahistidine tag. None of the parental complexes (i.e. antigen:fab complexes without any crystallization engineering applied to the Fab) produced crystals in the limited screens and time frame we employed (Table 3). All crystallization engineered complexes produced crystals, from 4 conditions (out of 192) for the G6 complex to 87 conditions for the H4 complex. Four of the six complexes produced structures, mostly at lower resolution. GITR complex crystals proved to be Fab alone upon solving the structure and TIGIT complex crystals did not diffract sufficiently well to produce a dataset. The crystals for both dupilumab and secukinumab required optimization to reach their respective 3 and 3.2Å resolutions (Fig 4A & B). Crystals from the initial screens diffracted to 5Å in the case of the former and not at all for the later.

**Table 3.** Crystallization of Fab:Antigen complexes. Complexes were screened in two 96-well crystallization screens and scored after 9 days. Number of conditions with crystals of any kind from the parental complex (without crystallization mutations) are indicated in the “Parent Xtal” column. Conditions with crystals from the CK1.0 Fab antigen complex in the column labelled “CK Xtal”. The best diffraction or dataset from these screens or subsequent optimizations is indicated in the “Res” column.

| Fab | Antigen | Parent Xtal | CK Xtal | Res (Å) | Notes | Reference |
| --- | --- | --- | --- | --- | --- | --- |
| G6 | Receptor | 0 | 4 | 3.6 | CK1.5 | Lilly molecule |
| Dupilumab | IL4Ra | 0 | 34 | 3.0 | optimized | KEGG D10354 |
| Secukinumab | IL17a | 0 | 54 | 3.2 | optimized | KEGG D09967 |
| h2155 | GITR | 0 | 54 | 2.6 | Fab only | US Patent 2013108641 |
| h22G2 | TIGIT | 0 | 30 | 9 |  | US Patent 2016176963 |
| H4 | Receptor | 0 | 87 | 2.6 |  | Lilly molecule |

**Fig 4.**
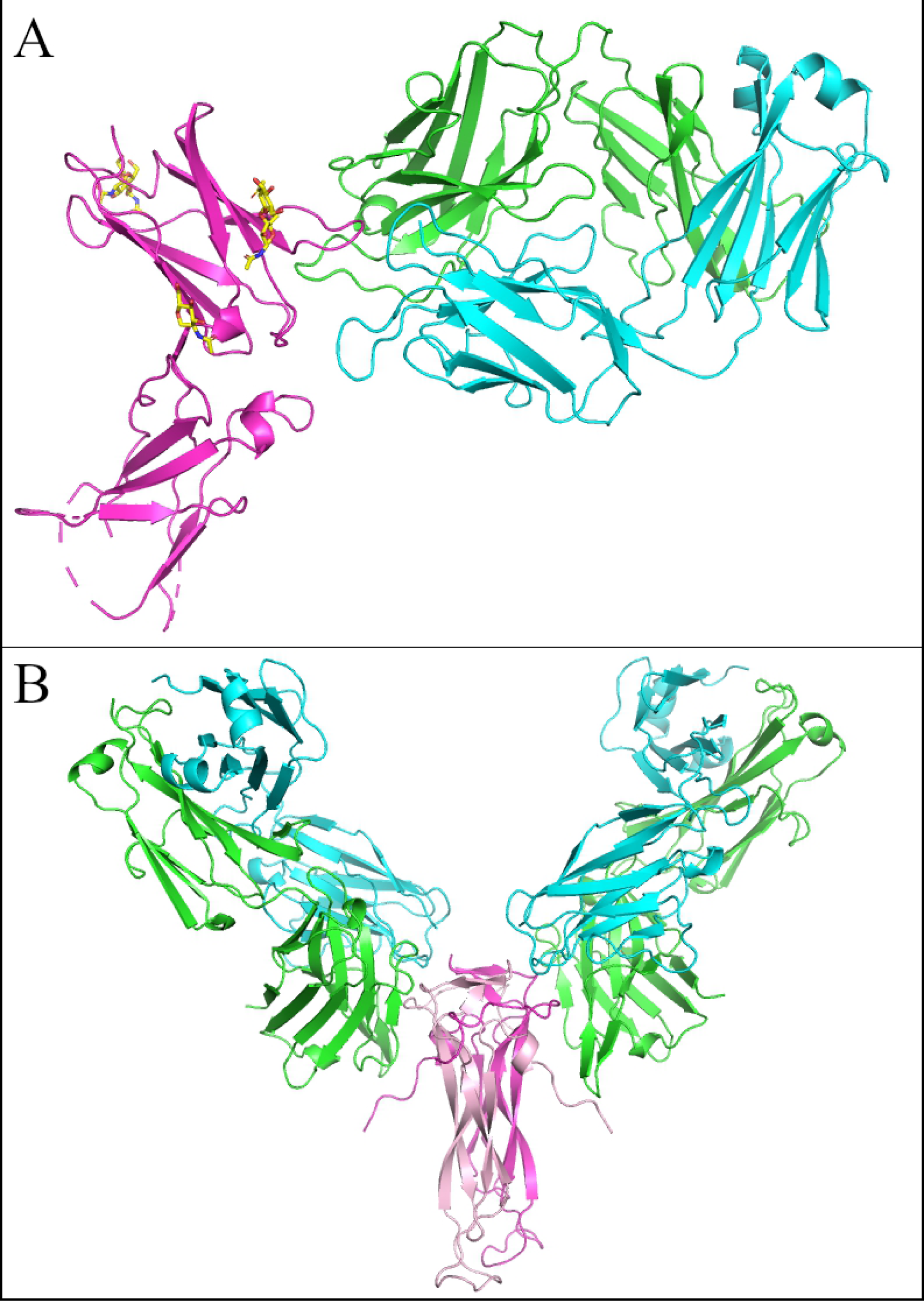
Crystal structures of Crystal Kappa variants. (A) Dupilumab Fab (HC in green and LC in cyan) complexed with human IL4R extracellular domain (in magenta and glycosylation in yellow sticks), (B) human IL17 dimer (magenta and pink) complexed with Secukinumab Fabs (HCs in green and LCs in cyan)

Because the secukinumab:IL17 complex produced low resolution but diffracting crystals, it was selected for further comparisons of CH1 domain isotype and C-terminus. Four new constructs were created: an IgG4 version that was 5 Residues shorter and ended with the sequence DKRVESK (“tagless”), one that ended DKRVH_6_ (“tagged”) and an IgG1 version ending with the sequence KSC with or without a hexahistidine tag. These were purified and screened as before at 10 mg/ml and also at 5 mg/ml. The shorter IgG4 versions produced fewer crystals than the original (24 for the tagged and 18 for the untagged versus 54 for the original tagged version). The IgG1 versions gave a similar number of conditions with crystals (88 tagged and 43 untagged versus 54 for the original). The 10 mg/ml IgG4 tagged version diffracted to 4.2Å, like the parent that didn’t initially diffract but produced 3.2Å after optimization. The IgG1 tagged version on the other hand at 10 mg/ml produced a 2.7Å dataset directly from the initial screen and the IgG1 untagged at 5 mg/ml produced a 2.4Å dataset directly from the initial screen.

### Column fraction crystallization

The robust crystallization of the engineered Fabs encouraged us to attempt crystallization directly from column fractions. The ΔQGTTSΔ variant of G6 was tested to determine the viability of this application (Fig 5). Column fraction crystallization produced the same crystal form as the purified and concentrated sample at 10 mg/ml. The CFC structure was at a respectable resolution of 2.4Å (versus 1.4Å).

**Fig 5.**
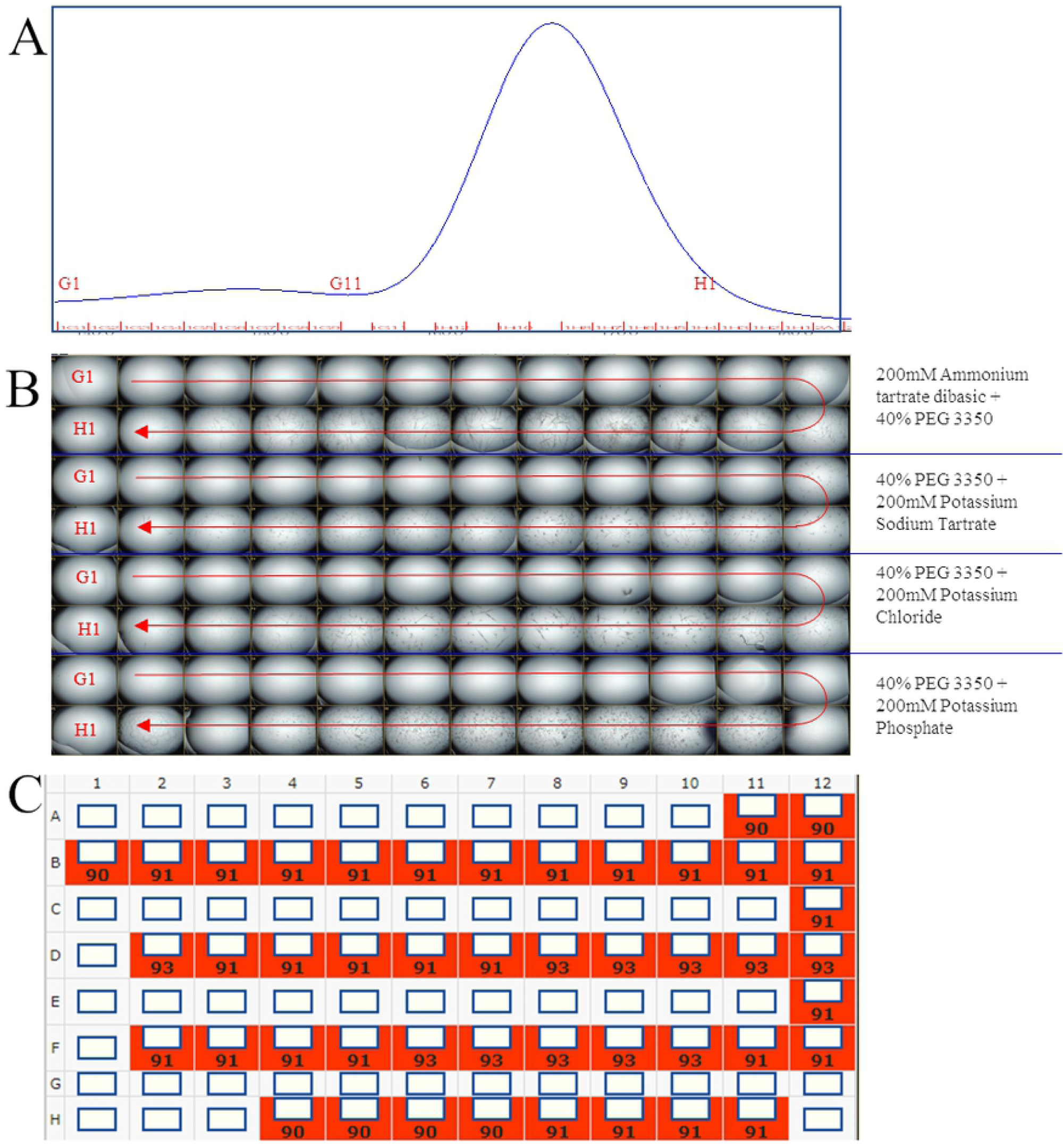
Column fraction crystallization. Column fractions from size exclusion chromatography were used directly in a vapor diffusion crystallization experiment in 4 conditions. (A) shows the chromatogram. (B) shows the images of every crystallization drop at day 9 (C) shows the score assigned to each, where anything crystalline is scored at 90 or higher

## Discussion

Crystallization of Fabs tends to be easier than most proteins. Because of this, Fabs are used as crystallization chaperones for membrane protein crystallization and other recalcitrant targets [11]. Still, relative to Fabs there are individual proteins and whole classes of proteins that crystallize faster, in simpler conditions, from heterogeneous mixtures, at very low concentrations, or a combination of these. In our decades of crystallizing proteins we have seen many that crystallize in just a few hours. We have seen crystals grow during protein concentration with no precipitant present. We have had a host cell protein undetectable in our target sample by-SDS-PAGE or mass spectrometry (*e. coli* carbonic anhydrase) crystallize sufficiently well to collect a better than 2.0Å dataset while attempting to crystallize the highly pure and concentrated target protein. The first antibody fragment structures from the 1970s were lambda light chain dimers (Bence-Jones proteins) extracted from urine and presumably heterogeneous by modern crystallization standards but able to crystallize [12]. Consider the name of the Fc portion of antibodies. It means “crystallizable fragment” and comes from the fact that after papain cleavage this fraction can be readily crystallized by dialysis against water [13].

Our goal was to create a universal Fab design that crystallizes with as many desired properties as possible, i.e. crystallize quickly, to a high resolution, from heterogeneous mixtures, at low concentrations, in many conditions. A design that incorporates only constant domain modifications is preferable as it would avoid complications due to germline differences in the variable domains as well as simplify cloning. We expected that such a Fab would also improve the crystallization of Fab:antigen complexes.

One of the difficult aspects of evaluating new techniques in crystallography is the fact that subtle changes can influence the nucleation and growth of protein crystals. While numerous techniques have been published and become part of the “toolkit” of crystallographers, their success rates are not known nor is the influence of other factors. To reduce this uncertainty we applied our designs to several targets, utilized comparable isotypes and C-termini, used the same purification, crystallization and crystal harvesting procedures (in parallel as much as possible), utilized crystal seeding to reduce variability in nucleation, evaluated crystallization experiments at the same elapsed time, and had the same scientists conduct the purification and crystallization across all experiments.

In a design counter to the natural tendency of proteins to avoid edge-to-edge aggregation, we designed constant kappa domain with a greater propensity to form edge-to-edge interactions [14]. The human Cκ variant domain (ΔQGTTSΔ) improved the frequency of crystallization by 50-fold for human Fabs. G6 parental Fab (fully human) yielded crystals in only one condition but the G6 that contained ΔQGTTSΔ yielded crystals in approximately 100 conditions (i.e. more than half of the 192 conditions). Dupilumab parental Fab yielded crystals in 4 conditions but dupilumab Fab that contained ΔQGTTSΔ yielded crystals in approximately 120 conditions. The modified FG loop of the Cκ domain (“Crystallizable Kappa” or more simply “Crystal Kappa”) also enabled crystallization of Fab:antigen complexes. The fold improvement cannot be calculated because none yielded crystals without the Crystal Kappa design. Most Crystal Kappa versions of the complexes yielded between 30 and 90 crystalizing conditions. It should be noted that these are statistics from two plates at one relatively conservative time point.

Unlike the crystallization behavior, we do not believe that firm conclusions can be drawn from the diffraction quality of the crystals. Harvesting, cryoprotecting and collecting from crystals is so variable that it would require harvesting dozens of crystals from dozens of conditions to be confident of conclusions. The diffraction of G6 might have been improved with the designs (1.2-1.7 vs. 3.4Å). Dupilumab Fab appeared to diffract approximately the same (1.4-2.2 vs. 2.0Å).

Interestingly one of the G6 crystals produced a dataset and structure with higher resolution (1.15Å) than was available for any Fab in the PDB (or in our own database) at the time, directly from one of two initial trays. At the time this suggested to us that the K126A mutation and disulfide removal might have an advantage, but subsequent results with dupilumab Fab indicated that the simplest ΔQGTTSΔ version was as good as the more complicated versions.

Relative to the Fab results, results with the complexes were somewhat disappointing. While they all produced crystals, a significant advantage to any effort, only 4 out of 6 yielded complex datasets good enough to solve and refine, and these tended towards lower resolutions. One (h2155 + GITR) produced crystals of the Fab alone.

The 3.0Å structure of Dupilumab Fab complexed with human IL4R shows an epitope that substantially overlaps with IL4 and IL13 binding, explaining its blocking activity (Fig 4A). The central part of the epitope is the CD loop [15], explaining why Dupilumab has no cross-reactivity with cynomolgous monkey IL4R which has a very different sequence in this region (L_67_L_68_ vs. Q_67_S_68_). The 3.0Å Secukinumab IL17 complex (and its 2.4Å improved structure) shows two Fabs bound to the IL17 dimer with a discontinuous epitope, each Fab binding portions of both IL17 chains (Fig 4B). The H4 complex produced a curious crystal packing arrangement with 3 Fabs in the asymmetric unit (not shown). One of the Fabs formed two HC:LC beta packing interactions typical for the CK design. Another Fab formed one on the LC side and nothing on the HC side. And the third Fab formed a HC:LC interaction with the second and its LC formed a LC:LC beta packing interaction, the only such interaction we have seen.

An additional refinement of the secukinumab constructs shortened the C-terminus of the IgG4 construct and included IgG1 versions for the first time and compared His_6_ tagged versus untagged versions. We also compared 10 mg/ml to 5 mg/ml in an attempt to reduce the crystal crowding for harvesting purposes. In this series, the IgG1 versions behaved better than the IgG4 and yielded a 2.7Å dataset for the tagged (at 10mg/ml) and 2.4Å dataset for the untagged Fab (at 5 mg/ml) versions directly from initials. This resolution improvement is probably real as the 3.0Å dataset from the CK1.0 construct was obtained after optimization and screening of numerous crystals. Interestingly the G1, G4 (two versions), tagged and tagless all produced isomorphous crystals. We do not yet know how general this result will be but believe optimizing the isotype and C-terminal His tag to be a fruitful avenue of further improvement, especially for complexes.

With regard to speed of crystallization, all the crystals described grew within a week and for those checked more frequently crystals appeared within hours. With regard to concentration, the column fraction crystallization experiment shows that it is possible to obtain crystals from samples as dilute as 0.1 mg/ml, at least for Fabs alone. Besides the implications for required concentrations, the CFC has other potential advantages. For example, the character of the protein at the leading edge of a column peak is probably different from the trailing edge and one or the other might be more productive in determining a structure. The CFC experiment and the fact that Fabs crystallize in more than half of crystallization conditions at high concentrations suggest that for Fabs the Crystal Kappa design should allow for a dramatically simplified set of screening conditions.

The rabbit Fabs form beta sheet crystal packing interactions mostly from LC to LC, with some HC to HC, and a few HC to LC. So we were surprised that in every crystal form of every Fab and Fab complex (including a dozen more not reported here) we saw only HC to LC constant domain packing (in two cases, no beta sheet packing). This produces a continuous thread of constant domains the entire length of the crystals. HC to HC beta packing is not common in human Fabs, but the absence of LC to LC packing with the Crystal Kappa design is curious.

The lambda light chain domains can also form the kind of beta sheet packing discussed here as either Fabs (e.g. 3tv3.pdb) or as Bence-Jones light chain dimers (e.g. 1dcl.pdb) [16]. Like the rabbit kappa, they have an FG loop that is shorter by two residues and can form beta sheets across crystal packing interfaces. Edmundson & Borrebaeck examined this interface in detail and highlighted the importance of “packing triads,” three alternating residues with a propensity to form beta sheets. Subsequent work focused on scFv crystallization [17]. Our work shows that the Crystal Kappa design of the FG loop (ΔQGTTSΔ) has the most dramatic influence on human Fab crystallization.

We believe we have created a crystallizable variant of the human constant kappa domain that dramatically improves the frequency of crystal formation for Fabs and Fab complexes, yields high resolution structures for Fabs (and Fab:peptide complexes) and can yield in most cases at least low resolution datasets and structures of Fab:protein complexes. The Crystal Kappa design appears to allow for overnight crystallization from dilute samples in screens of a handful of conditions. Crystal Kappa designs should make Fab structure determination robust even using smaller screens and less protein and speed up complex structure determination, including Fab chaperone complexes with difficult targets.

## Materials and Methods

### Engineering and molecular biology

Analysis of rabbit Fab crystal packing was conducted in Pymol utilizing structures available in the Protein Databank and Eli Lilly’s proprietary structural database. Alignment of LC constant domains from various species utilized BLAST. Amino acid sequence for the variable domains of dupilumab and secukinumab were obtained from the Kyoto Encyclopedia of Genes and Genomes website [18], entries D10354 [19] and D09967 [20]. Sequences for h2155 and h22G2 were obtained from patents [21,22]. Expression vectors were created by synthesizing the corresponding DNA fragments as gblocks (IDT, Coralville IA) and cloning into mammalian expression vectors using standard techniques.

### Expression and Purification

Fabs and antigen ECDs were expressed in mammalian cell culture CHO cells. Protein containing cell culture supernatants were harvested and clarified media was purified by Immobilized Metal Affinity Chromatography (IMAC) using His Trap™ Excel (GE Healthcare) using PBS buffer plus 15mM Imidazole, pH 7.5 as the binding buffer. The proteins were then elution on a 10 column volume gradient elution in PBS plus 0.3M Imidazole, pH 7.4. The eluents were collected and concentrated using a Millipore 10 KDa spin concentrator. The concentrate IMAC pools were loaded on either a Superdex75 or Superdex200 (GE Healthcare) columns. Proteins were concentrated again to 10 mg/ml for crystallization trays.

The proteins were characterized by analytical size exclusion chromatography (Waters) and SDS-PAGE gel (S1 and S2 Figs).

### Crystallization and Structure Determination

All samples were concentrated to 5-10 mg/ml and were set up at room temperature in vapor diffusion sitting drops at a ratio of 1:1 using Qiagen Classics II and PEGs crystallization screens. Drops were immediately cross seeded with related Fab crystal seeds for the Fabs crystallization and complex-crystal seeds for complex crystallization. Images of crystallization trays were taken on day 1, day 4, and day 9. Prior to freezing in liquid nitrogen, crystals were transferred to a cryoprotectant solution consisting either of well solution supplemented with an additional 10% of the precipitant used in that crystallization well and 25% of glycerol or from the mother liquor, if it included a precipitant with cryoprotecting qualities (such as PEG 400, PEG MME 550, PEG MME 2K etc.) in concentrations sufficient for cryoprotection.

Structure determination diffraction datasets were collected at the following sources: Lilly Research Collaborative Access Team (LRL-CAT) Beamline 31-ID at Advanced Proton Source (Argonne, IL); Beamline ALS-502 at Advanced Light Source (Berkley, CA); Beamline I04-1 at Diamond Light Source (Oxfordshire, UK). Data were integrated and reduced using MOSFLM [23] and the CCP4 suite of programs [24]. Initial molecular replacement solutions were obtained using Phaser (CCP4 suite) [25]. The model was built using COOT [26] and refined using Refmac [27] or Buster [28] and validated using internal developed protocols.

## Abbreviations

CDR: Complementarity determining region
CFC: Column fraction crystallization
CK: Crystal Kappa
GITR: Glucocorticoid-Induced TNFR-Related Protein
TIGIT: T-cell immunoreceptor with Ig and ITIM domains
LC: Light Chain
HC: Heavy Chain
CH1: Constant Heavy Domain 1
Ckappa: Constant Kappa Domain
Fv: Antibody variable domain fragment

## Acknowledgments

Special thanks to Craig Dickinson, Stephen Demarest and Xiaomin Yang for many helpful discussions, to Jordi Benach and the beamline staff for data collection and processing, to Jorg Hendle for crystallographic quality control, to Nicole Niemela for assistance in small scale protein purification, and to the remaining staff of the Protein Biosciences, Crystallization, Crystallography, and Beamline groups for help in countless small ways. This research used resources of the Advanced Photon Source, a U.S. Department of Energy (DOE) Office of Science User Facility operated for the DOE Office of Science by Argonne National Laboratory under Contract No. DE-AC02-06CH11357. Use of the Lilly Research Laboratories Collaborative Access Team (LRL-CAT) beamline at Sector 31 of the Advanced Photon Source was provided by Eli Lilly Company, which operates the facility. This research used resources of the Advanced Light Source, which is a DOE Office of Science User Facility under contract no. DE-AC02-05CH11231. Beamline 5.0.2 of the Advanced Light Source, a DOE Office of Science User Facility under Contract No. DE-AC02-05CH11231, is supported in part by the ALS-ENABLE program funded by the National Institutes of Health, National Institute of General Medical Sciences, grant P30 GM124169-01. The authors would like to thank Diamond Light Source for beamtime, and the staff of beamlines I04 for assistance with crystal testing and data collection.

List of structures deposited in the Protein Databank are in S1 Table.

## Supporting Information

**S1 Table. Crystal structures deposited in the Protein Data Bank**.

**S1 Fig. Small scale expression/purification to eliminate variants for scale up**. Shows well-formed Fabs. A) Fabs are expression and IMAC purified. aSEC shows fabs are well formed. B) Non-reduced SDS-PAGE gel shows Fabs are intacted.

**S2 Fig. Large scale purification reproduced quality Fabs**

## References

1. Kaplon H, Muralidharan M, Schneider Z, Reichert JM. Antibodies to watch in 2020. MAbs. 2020 Jan-Dec;12(1):1703531. doi: 10.1080/19420862.2019.1703531

2. Lu RM, Hwang YC, Liu IJ, Lee CC, Tsai HZ, Li HJ, et al. Development of therapeutic antibodies for the treatment of diseases. J Biomed Sci. 2020 Jan 2;27(1):1. doi: 10.1186/s12929-019-0592-z

3. Parmley SF, Smith GP. Antibody-selectable filamentous fd phage vectors: affinity purification of target genes. Gene. 1988 Dec 20;73(2):305–18. doi: 10.1016/0378-1119(88)90495-7

4. Boder ET, Wittrup KD. Yeast surface display for screening combinatorial polypeptide libraries. Nat Biotechnol. 1997 Jun;15(6):553–7. doi: 10.1038/nbt0697-553

5. Lonberg N. Human monoclonal antibodies from transgenic mice. Handb Exp Pharmacol. 2008;(181):69–97. doi: 10.1007/978-3-540-73259-4_4

6. Carter PJ, Lazar GA. Next generation antibody drugs: pursuit of the ‘high-hanging fruit’. Nat Rev Drug Discov. 2018 Mar;17(3):197–223. doi: 10.1038/nrd.2017.227

7. Xu Y, Wang D, Mason B, Rossomando T, Li N, Liu D, et al. Structure, heterogeneity and developability assessment of therapeutic antibodies. MAbs. 2019 Feb/Mar;11(2):239–264. doi: 10.1080/19420862.2018.1553476

8. Slabinski L, Jaroszewski L, Rodrigues AP, Rychlewski L, Wilson IA, Lesley SA, et al. The challenge of protein structure determination--lessons from structural genomics. Protein Sci. 2007 Nov;16(11):2472–82. doi: 10.1110/ps.073037907

9. Terwilliger TC, Stuart D, Yokoyama S. Lessons from structural genomics. Annu Rev Biophys. 2009;38:371–83. doi: 10.1146/annurev.biophys.050708.133740

10. McPherson A. Crystallization of biological macromolecules. Cold Spring Harbor: Cold Spring Harbor Laboratory Press; 1999.

11. Griffin L, Lawson A. Antibody fragments as tools in crystallography. Clin Exp Immunol. 2011 Sep;165(3):285–91. doi: 10.1111/j.1365-2249.2011.04427.x

12. Wilson DW. A spontaneous crystallization of a bence-jones protein. J Biol Chem. 1923 56:203–214.

13. Porter RR. The hydrolysis of rabbit y-globulin and antibodies with crystalline papain. Biochem J. 1959 Sep;73(1):119–26. doi: 10.1042/bj0730119

14. Richardson JS, Richardson DC. Natural beta-sheet proteins use negative design to avoid edge-to-edge aggregation. Proc Natl Acad Sci U S A. 2002 Mar 5;99(5):2754–9. doi: 10.1073/pnas.052706099

15. Ul-Haq Z, Naz S, Mesaik MA. Interleukin-4 receptor signaling and its binding mechanism: A therapeutic insight from inhibitors tool box. Cytokine Growth Factor Rev. 2016 Dec;32:3–15. doi: 10.1016/j.cytogfr.2016.04.002

16. Edmundson AB, Borrebaeck CA. Progress in programming antibody fragments to crystallize. Immunotechnology. 1998 Jan;3(4):309–17. doi: 10.1016/s1380-2933(97)10002-1

17. Wingren C, Edmundson AB, Borrebaeck CA. Designing proteins to crystallize through beta-strand pairing. Protein Eng. 2003 Apr;16(4):255–64. doi: 10.1093/proeng/gzg038

18. Kanehisa M, Goto S. KEGG: kyoto encyclopedia of genes and genomes. Nucleic Acids Res. 2000 Jan 1;28(1):27–30. doi: 10.1093/nar/28.1.27

19. KEGG DRUG: Dupilumab [cited 2020 Apr 2]. Database: KEGG [Internet]. Available from: https://www.kegg.jp/dbget-bin/www_bget?dr:D10354

20. KEGG DRUG: Secukinumab [cited 2020 Apr 2]. Database: KEGG [Internet]. Available from: https://www.kegg.jp/dbget-bin/www_bget?dr:D09967

21. Baurin N, Blanche F, Cameron B, Dabdoubi T, Fordham J, Kominos D, et al., inventors; Sanofi SA assignee. Anti-gitr antibodies. United States patent US 2013108641. 2013 May 2.

22. Maurer MF, Chen TT, Devaux B, Srinivasan M, Julien SH, Sheppard PO, et al., inventors; Bristol Myers Squibb Co assignee. Antibodies to tigit. United States patent US 2016176963. 2016 Jun 23.

23. Leslie AGW & Powell HR. In: Read RJ, & Sussman JL (Eds.), Evolving methods for macromolecular crystallography: the structural path to the understanding of the mechanism of action of CBRN Agents. Dordrecht: Springer Netherlands; 2007

24. Winn MD, Ballard CC, Cowtan KD, Dodson EJ, Emsley P, Evans PR, et al. Overview of the CCP4 suite and current developments. Acta Crystallogr D Biol Crystallogr. 2011 Apr;67(Pt 4):235–42. doi: 10.1107/S0907444910045749

25. McCoy AJ, Grosse-Kunstleve RW, Adams PD, Winn MD, Storoni LC, Read RJ. Phaser crystallographic software. J Appl Crystallogr. 2007 Aug 1;40(Pt 4):658–674. doi: 10.1107/S0021889807021206

26. Emsley P, Lohkamp B, Scott WG, Cowtan K. Features and development of Coot. Acta Crystallogr D Biol Crystallogr. 2010 Apr;66(Pt 4):486–501. doi: 10.1107/S0907444910007493

27. Murshudov GN, Skubák P, Lebedev AA, Pannu NS, Steiner RA, Nicholls RA, Winn MD, Long F, Vagin AA. REFMAC5 for the refinement of macromolecular crystal structures. Acta Crystallogr D Biol Crystallogr. 2011 Apr;67(Pt 4):355–67. doi: 10.1107/S0907444911001314

28. Bricogne G. Direct phase determination by entropy maximization and likelihood ranking: status report and perspectives. Acta Crystallogr D Biol Crystallogr. 1993 Jan 1;49(Pt 1):37–60. doi: 10.1107/S0907444992010400

